# Anatomical insights into the vascular lay-out of the barley rachis: implications for transport and spikelet connection

**DOI:** 10.1101/2023.11.23.568454

**Authors:** Twan Rutten, Venkatasubbu Thirulogachandar, Yongyu Huang, Nandhakumar Shanmugaraj, Ravi Koppolu, Stefan Ortleb, Götz Hensel, Jochen Kumlehn, Michael Melzer, Thorsten Schnurbusch

## Abstract

**Background and aims:** Vascular patterning is intimately related to plant form and function. However, morphologic a l studies on the vascular anatomy of cereal crops, and inflorescences in particular, are scarce despite their importance for grain yield determination. Here, using barley (*Hordeum vulgare*) as a model, we study the vascular anatomy of the spike-type inflorescence. Our goal is to clarify the relationship between rachis (spike axis) vasculature and spike size, the implications for transport capacity and its interaction with the spikelets.

**Methods:** We employed serial transversal internode sections in multiple barley lines with different spike size, and investigated the internode diameter, vascular area and vein number size along the mature barley rachis. We then modeled the vascular dynamics along the main spike axis, and analyzed their relationship with spike size.

**Key results:** Internode diameter and total vascular area have a clear positive correlation with spike size whereas vascular number is only weakly correlated. While the lateral periphery of the rachis contains large mature veins of constant diameter the central part is occupied by a staggered array of small immature veins. This underlines the importance of minimizing transport resistance and suggests that transport and distribution of nutrients are spatially separated. Spikelet-derived veins enter the rachis either in the central area, where they often merge with the immature rachis veins, or in the periphery where they do not merge with the large mature veins. An increase in floret fertility through the conversion of a two-rowed barley into an isogenic six-rowed line, as well a decrease in floret fertility due to enhanced pre-anthesis tip degeneration caused by the mutation *tip sterile 2.b* (*tst2.b*) significantly affected vein size, but had limited to no effects on vein number or rachis diameter. Comparative analysis of a wild barley accession suggests that the domestication of barley may have favored plants with enhanced rachis transport capacity.

**Conclusions:** The rachis vasculature is the result of a two-step process involving an initial lay-out followed by size adjustment according to floret fertility/spike size. The functional processes of long distance transport and local supply to spikelets are spatially separated while a vascular continuity between rachis and spikelets appears non-essential.

## Introduction

In cereals, the rachis’s vasculature varies considerably from that of the shoot. Already Percival (1921) stressed that while in wheat (*Triticum aestivum*), the shoot displays a consistent shape and vascular organization throughout, the corresponding features of the rachis change from semicircular to spindle-shaped within several internodes. The same author also suggested an acropetal decline in vascular number within the rachis. In barley (*Hordeum vulgare*), Kirby and Rymer (1975) discerned three types of vascular bundles comprising of two opposite crescent rows of median veins demarcated by two large lateral veins, the whole of it flanked by a variable number of small peripheral veins. They further reported that at each rachis node, the two lateral veins and the whole arch of median veins adjacent to the spikelet attachment site divides, with one branch entering the spikelets and the other proceeding along the rachis. The median veins opposite the spikelets, however, do not divide but continue to the next rachis node. Studying only spike fragments, Kirby and Rymer (1975) refrained from explicitly commenting on an acropetal decline in vascular bundles. Whingwiri et al. (1981) confirmed the existence of a gradual acropetal decline in both vein number and vein size in wheat spikes. Similar to Evans et al. (1970), they noted a striking near 1:1 relationship between the number of spikelets and the number of main vascular bundles at the base of the rachis. The idea of each spikelet being assigned to a singular vascular bundle, first expressed by Evans et al. (1970), was slight ly revised into a model in which each spikelet is supplied by a constant number of one or two bundles (Whingwiri et al., 1981). In line with this, Bremner (1972) concluded that wheat spikelets probably have their supporting vascular bundles linked in parallel, whereas the kernels within the spikelet are connected in series.

The simplicity of the “one vein for one spikelet” model is appealing but hardly applicable to barley, in which the number of spikelets can easily exceed the number of rachis veins by a factor of three or more. A gradual acropetal decline in vein size, as reported for wheat, also has profound physical implications, as it will greatly increase the resistance to assimilate transport towards the apex (Chiavarria and Pessoa dos Santos, 2012). The phenomenon of pre-anthesis tip degeneration in barley spikes and wheat spikelets, which is the failure of the most apical spikelets/florets to develop into grains (Appleyard et al., 1982; Alqudah and Schnurbusch, 2014; Boussora et al., 2019; Thirulogachandar and Schnurbusch, 2021), has sometimes been suggested to be a result of transport limitations caused by increased resistance (Kirby and Faris 1970, Hanif and Langer 1972, Whingwiri and Kemp 1980). Nevertheless, when Bremner and Rawson (1978) removed the basal grains of wheat spikelets, they observed an improved growth rate of the more distal grains suggesting that the limiting factor is not transport capacity but the availability of resources. Subsequent spikelet and grain removal experiments in rice (Kato, 2004, You et al., 2016) and wheat (Ma et al., 1996) confirmed these observations. Whingw ir i et al. (1981) considered it premature to judge the vasculature of the spike to be a static entity. In wheat, Evans et al. (1970) noticed a clear correlation between spikelet number and both size and number of vascular strands within the peduncle. This indicates the presence of a dynamic component in vascular development, which, according to Whingwiri et al. (1981), probably exerts itself in the early stages of spike development. Waddington et al. (1983) suggested the existence of two distinct dynamic components. The first being the developmental progress that ends when maximum yield potential is reached (Thirulogachandar and Schnurbusch, 2021). Beyond this follows a phase of exponential growth of the initiated organs whose fates are determined before anthesis. Accordingly, the lay-out of the main rachis vasculature should be completed by the time of maximum yield potential and its final dimensions fixed before the start of anthesis.

Trying to shed more light on the vascular organization of the barley rachis, we investiga ted parameters including rachis diameter, number and size of the main vascular bundles as well as their distribution within serial transversal internode sections obtained from mature spikes of the two-rowed cv. Golden promise (two-rowed GP). The influence of spikelet fertility on rachis vasculature was studied in an isogenic six-rowed GP*-vrs1* line generated by targeted mutagenesis (Thirulogachandar et al., 2023), while for the effect of a reduction in fertile spikelets, we compared the cv. Bowman with its near-isogenic mutant BW*-tst2.b* (Huang et al., 2023) exhibiting extended pre-anthesis tip degeneration. Our results confirm the assumption of Waddington et al (1983) that the morphogenesis of the barley rachis vasculature is the result of two distinct processes: initiation and processing, with ultimate vein size strongly correlating to the number of fertile spikelets. While mature veins of constant size are located in the rachis periphery, immature veins of declining size occupy the middle part, suggesting a spatial separation between long distance transport and local delivery to the spikelets. Our data further indicate that, while spikelet-derived veins can enter the rachis both in the outer periphery and in the central part, they are only capable of fusing with the immature rachis veins residing in the center. A comparative analysis of the wild barley accession HID003 suggests that barley domestication may also have favored a selection for plants with improved rachis transport capacity.

### 2.1. Plant material

The two-rowed barley (*Hordeum vulgare* L.) cvs. Bowman and Golden Promise (two-rowed GP) as well as wild barley (*H. vulgare* L.*, subsp. spontaneum* (K. Koch)) accession HID003 were obtained from the IPK gene bank. The barley mutant, *tst2.b* (BW883) was ordered from NordGen seed bank. To reduce background introgressions from the original mutagenesis recipient (cv. Donaria), we further backcrossed BW883 to wild-type Bowman to generate BC6F4∼5 plants (Huang et al., 2023). The isogenic six-rowed GP*-vrs1* line was obtained through targeted mutagenesis conducted by CRISPR-associated (Cas) endonuclease technology (Thirulogachandar et al., 2023). A guide RNA (TCTGGAGCTGAGCTTCCGGGAGG) was designed for the homeodomain region of the *Vrs1* gene and inserted between the OsU3 promoter and the downstream gRNA scaffold present in a generic, monocot-compat ib le intermediate vector pSH91 (Budhagatapalli et al., 2016). Next, the whole expression cassette of the gRNA-Cas9 was introduced into the SfiI cloning site of the binary vector p6i-d35S-TE9 (DNA-Cloning-Service, Hamburg, Germany). Transgenic plants were created in the two-rowed cv. Golden Promise, via Agrobacterium-mediated table transformation (Hensel et al., 2009). The two-rowed GP and its gene-edited six-rowed GP*-vrs1* were grown in 14 cm pots, cv. Bowman, line BW*-tst2.b* and wild barley acc. HID003 were grown in 9 cm pots. Plants were cultivated under greenhouse conditions at 22/18°C in 16 h/8 h days-night.

### 2.2. Rachis vasculature measurements

Spikes were collected after anthesis, by which time development and growth of the barley spike, including its vasculature, is fully completed (Waddington et al., 1981, Yang & Zhang, 2006). Spikes were usually collected from prime tillers (see Table 1 for overview of plant material). In case of two-rowed GP mature spikes were also collected from secondary tillers to obtain a larger variation in spike size. Starting with the peduncle, which was sampled two cm underneath the base of the rachis, and ending with the first internode of the zone of pre-anthesis tip degeneration, free hand transverse sections were made at about one-fourth of the rachis internodes. Sections were examined in a Zeiss LSM780 laser scanning microscope (Carl Zeiss, Jena, Germany) using a 10x NA 0.45 objective (zoom 1, image size 1024 x 1024 pixels combined with tiling and Z-stacks with ensuing maximum intensity projection). Probes were scanned with a 405 nm laser line and cell wall autofluorescence recorded with a 406-500 nm bandpass. For area measurements the open source Fiji software (Schindelin et al., 2012) was used. Surface area of individual veins was determined as the area surrounded by a bundle sheath. Classical Student’s t-tests were performed to reveal significant differences between data sets.

**Table 1.**
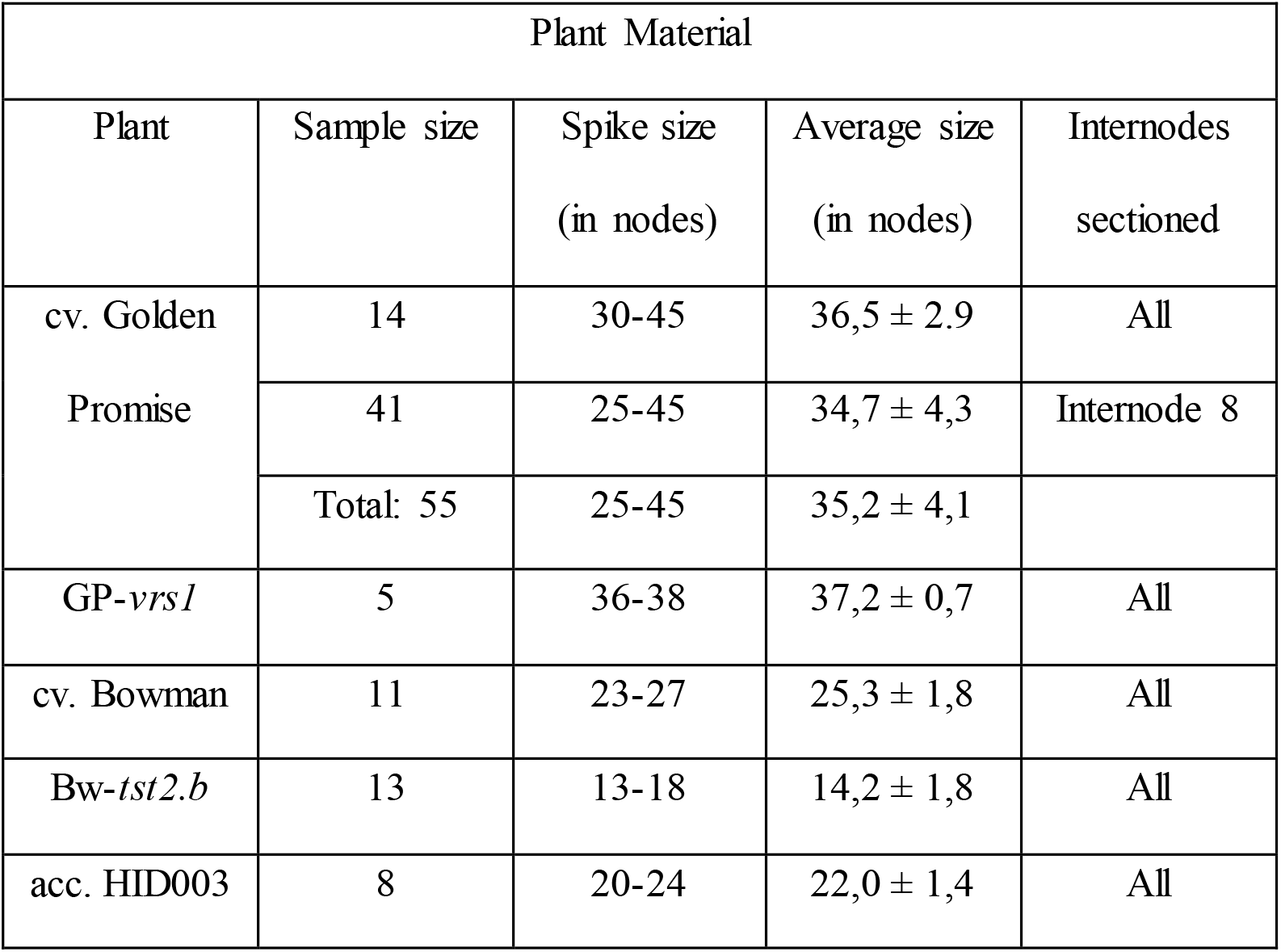
Overview of plant material used.

## Results

### Basic morphology of the barley rachis

To create a reference model for the shape and vascular organization of the barley rachis, we used fully differentiated spikes of two-rowed GP (Fig. 1a), with node number varying between 30 to 43 (average 36,5 ± 2,9 nodes, n = 14). By employing serial transversal internode sectioning, each vein on its course between two nodes is cut both shortly after and shortly before accessing a node (Fig. 1b). The persistent morphological features of individual veins allow them to be followed throughout the rachis. The latter starts as a semicircular cylindrical structure that rapidly attains an ellipsoid shape [**supplementary information – video**]. The peduncle at the base of the rachis displays two rings of vascular bundles of which the outer ring of smaller bundles gives rise to the outer lateral veins mentioned by Kirby and Rymer (1974) (Fig. 1c). The inner ring of larger bundles develops into the main rachis vasculature consisting of two opposing crescent rows of median veins with two large lateral veins at the vertices (Fig. 1c) [**supplementary information – video**]. Due to their different origin and different morphological features, including much smaller metaxylem vessels, the outer lateral bundles were excluded from the present study which thus deals exclusively with the median and lateral veins (Fig. 1e). Unless stated otherwise the data presented here was collected from 14 mature spikes of two-rowed GP in the late stages of grain filling (average size 36,5 ± 2,9 nodes, n = 14).

**Fig. 1.**
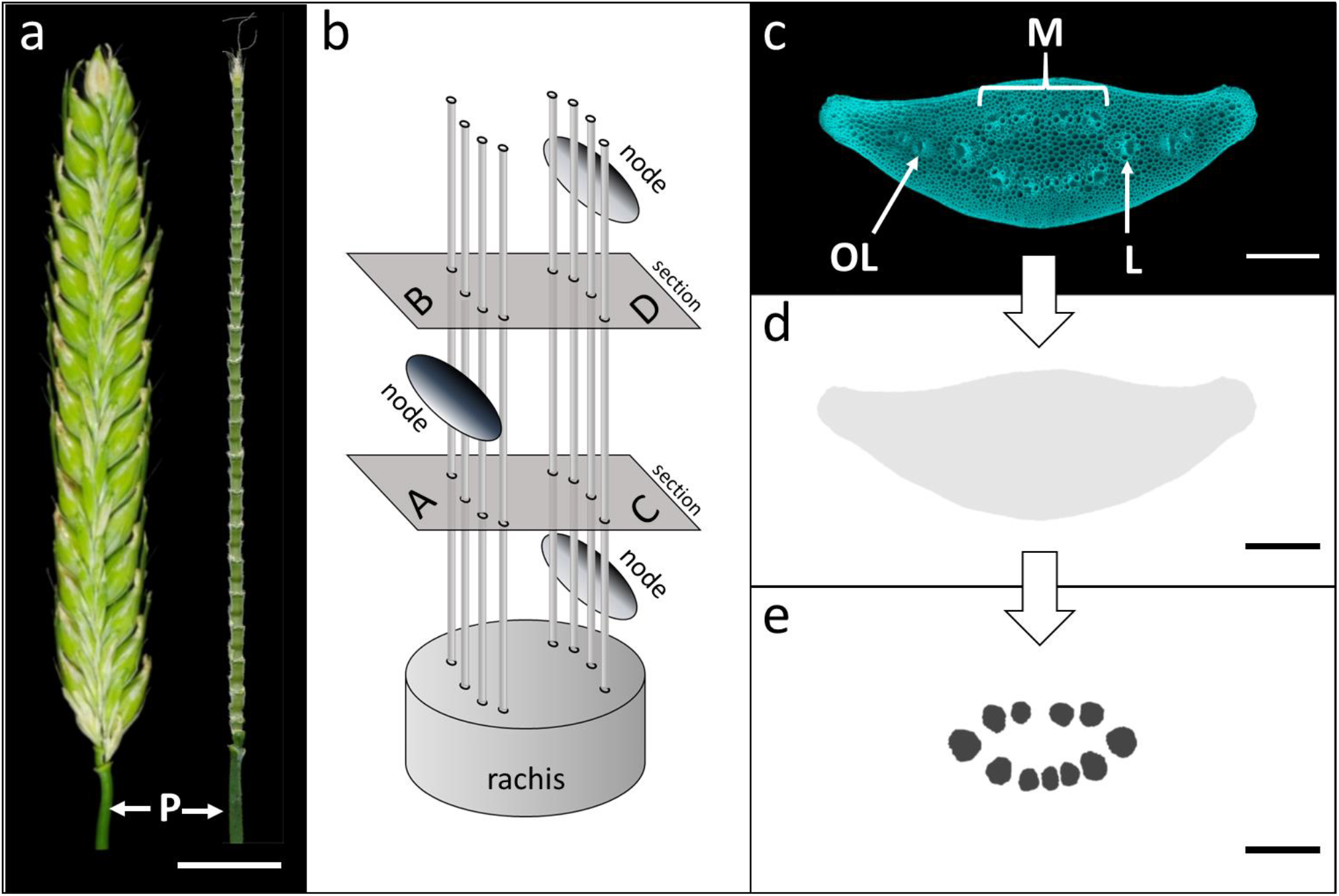
Barley cv. Golden Promise. (a) Mature Spike and central view of rachis after removal of spikelets. P = peduncle. (b) Schematic view of rachis showing the two opposite rows of median veins and the alternate arrangement of nodes on either side of the rachis. Serial internode sectioning enables the analysis of vascular changes happening after passing a node (A to B = internodal) and between two nodes (C to D = intranodal). (c) Transverse internode section reveals three types of vascular bundles: median (M), lateral (L) and outer lateral (OL). (d) Schematic view of internode diameter in grey. (e) Schematic view of size and distribut ion of main vascular bundles. Bar = 1000 µm (a), 200 µm (c, d, e).

### Rachis parameters

For a reconstruction of the vascular organization within the rachis we first focused on the parameters internode diameter, total vascular area and number of vascular bundles. Internode diameter reaches a weak local maximum at internode 3 and then decreases subsequently (Fig. 2a). The fall off is sharp before internode 10 but becomes gentler afterwards. The dynamic of the parameter total vascular area is remarkably similar to that of the internode diameter (compare Fig 2a and 2b) including a small local maximum between internode 2 and 3, followed by a decline which is steep until about internode 10 and much more gradual thereafter. The third parameter, vein number, reaches an absolute peak at internode 3 or 4 before gradually declining towards the apex (Fig. 2c).

**Fig. 2.**
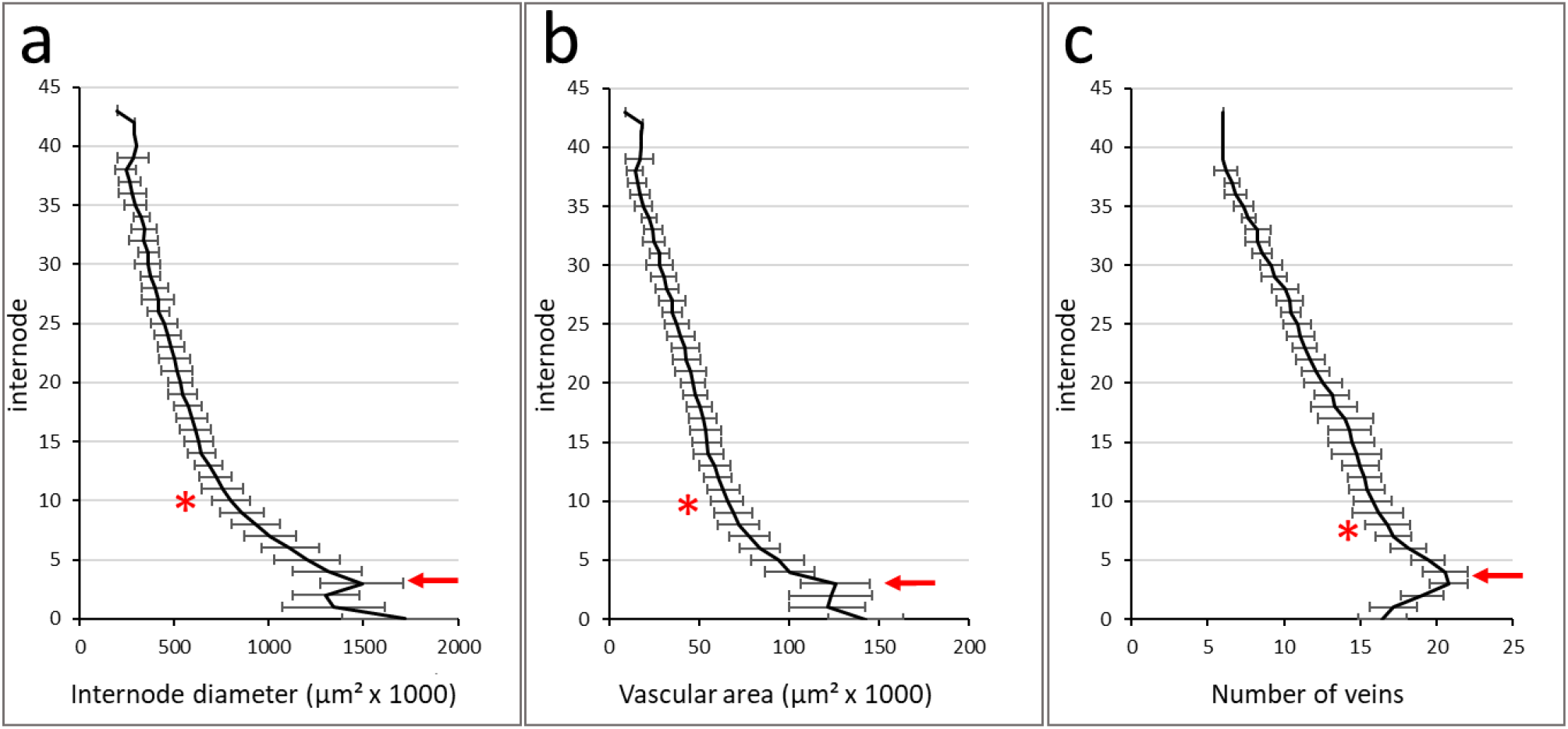
Dynamics of internode parameters along the rachis. (a) Internode diameter and (b) total vascular area reach a local maximum at internode 3 (red arrow) followed by a steep decline until around internode 10 (asterix) and a much more gradual decline thereafter. (c) Number of veins peaks near internode 4 (arrow) before gradually declining towards the apex.

### Rachis parameters and spike size

To determine the relationship between rachis parameters and spike size, a total of 55 spikes of two-rowed GP with node number varying between 25 and 45 were used (average size 35,2 ± 4,1 nodes). From each of these spikes internode 8 was collected and internode diameter, total vascular area, total lateral vein area and the area of the metaxylem vessels of the lateral veins (Fig. 3a-b) were measured.

**Fig. 3.**
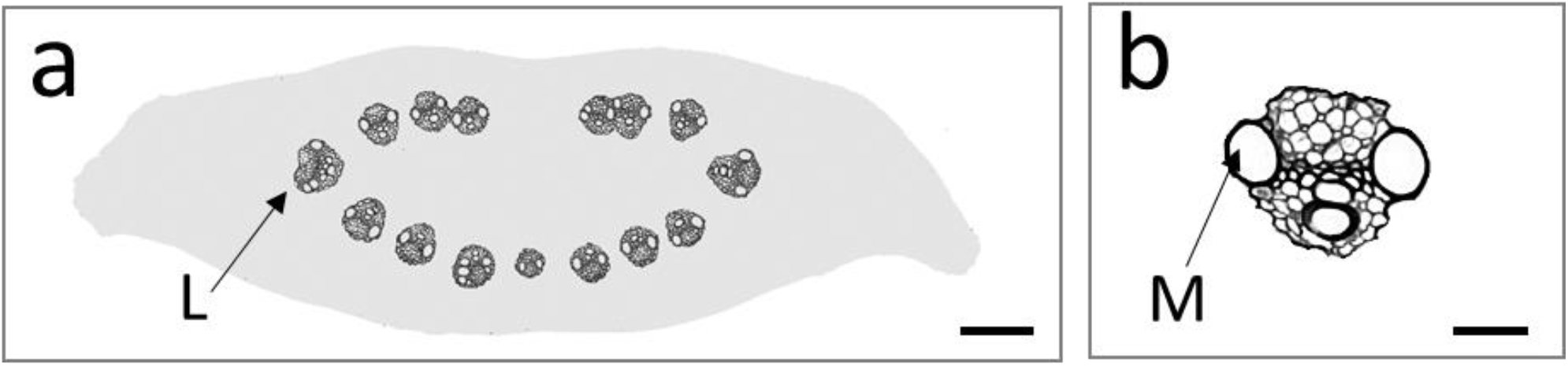
Schematic illustration of diameter and vascular content of internode no. 8 from a 36-node spike (a) with lateral vein in detail (b). L = lateral vein, M = metaxylem vessel. Bar = 100 µm (a), 20 µm (b).

The results, presented as dot plots, demonstrate that internode diameter (Fig. 4a) and total vascular area (Fig. 4b) show a clear positive correlation to spike size. In contrast to this, vein number only displays a weak positive correlation (Fig. 4c). The same data was then used, to investigate the relationships among the vascular parameter themselves. The outcome indicates that within a given internode there are fixed ratios between internode diameter and the vascular parameters total vascular area (Fig. 5a), individual vein size (Fig. 5b) down to metaxylem vessel size (Fig. 5c). Since these ratios are thus independent of spike size, a feature like internode diameter can be used to compare relative differences in transport capacities between spikes of different sizes.

**Fig. 4.**
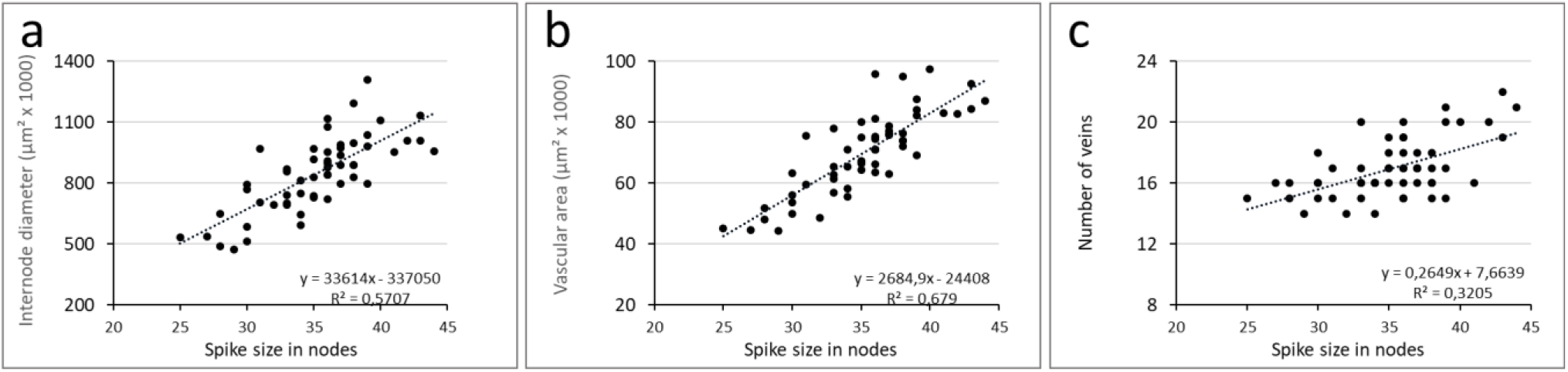
Rachis parameters in relation to spike size. Data was collected for internode no. 8 from a total of 55 spikes (average size 35,2 ± 4,1 nodes). Internode diameter (a) and total vascular area (b) show a clear positive correlation to spike size whereas vein number (c) shows only a weak positive correlation.

**Fig. 5.**
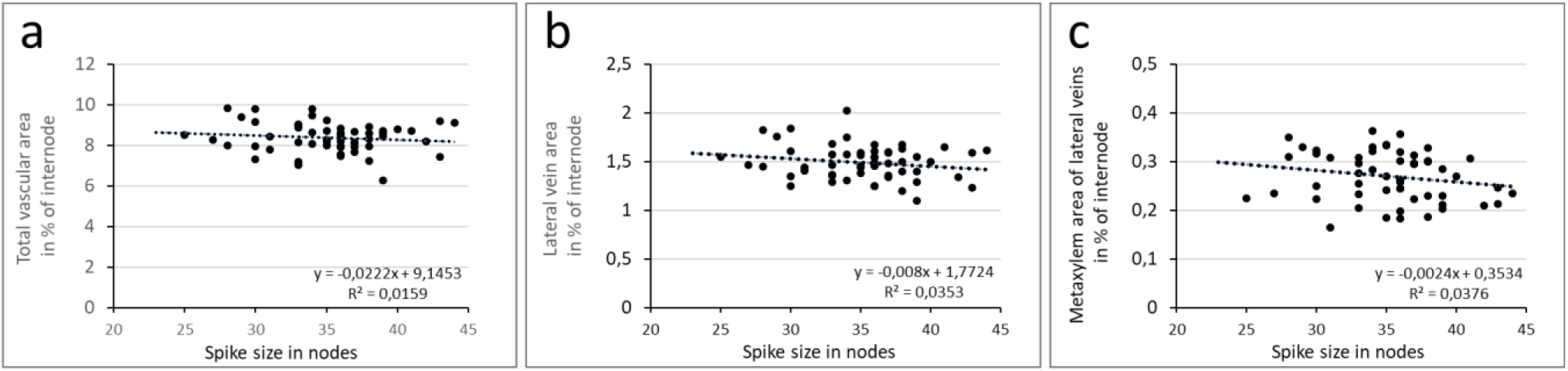
Vascular parameters in relation to internode diameter. Data was collected for internode no. 8 from a total of 55 spikes (average size 35,2 ± 4,1 nodes). Total vascular area (a), total lateral vein area (b) and total area of metaxylem vessels from the lateral veins (c) as percentage of internode diameter in relation to spike size. The results indicate that the ratios between internode diameter and vascular parameters down to metaxylem vessels are independent of spike size.

### Dynamics of individual vein orders

Treating the rachis vasculature as a whole, masks the behavior of its individual components. This is exemplified by a differentiated representation of the vascular bundles in a single 43-node rachis (Fig. 6). The graph for the total main vascular surface area shows the characteristic peak at internode 3, a sharp decline until about internode 10, and concluding smooth decline thereafter (Fig. 6a). To distinguish the contributions of individual veins, the median vasculature was divided into veins of ascending order (Fig. 6b). Here we followed the proposition of Kirby and Rymer (1975) that consecutive new classes of median veins develop inward of pre-existing ones. Inwards of the lateral veins thus follow several orders of median veins each comprising of four vascular bundles (Fig. 6b). For clarity all veins beyond third order were grouped as fourth order veins. The results show that all vein orders initially mimic the dynamics of the total vascular area in showing a peak value around internode 2 followed by a brief steep decline (Fig. 6c, suppl. Fig. 2). While the vascular area of 4^th^ order veins displays a continuous decline and vanishes around internode 15, the other vein orders show increasingly long zones of size stability and do not start their final decline before the preceding vein order has disappeared (Fig. 6c, d). Accordingly, the middle part of the rachis is thus occupied by a staggered array of declining, probably immature veins while stable sized, probably mature, veins occupy the lateral regions (Fig. 6d).

**Fig. 6.**
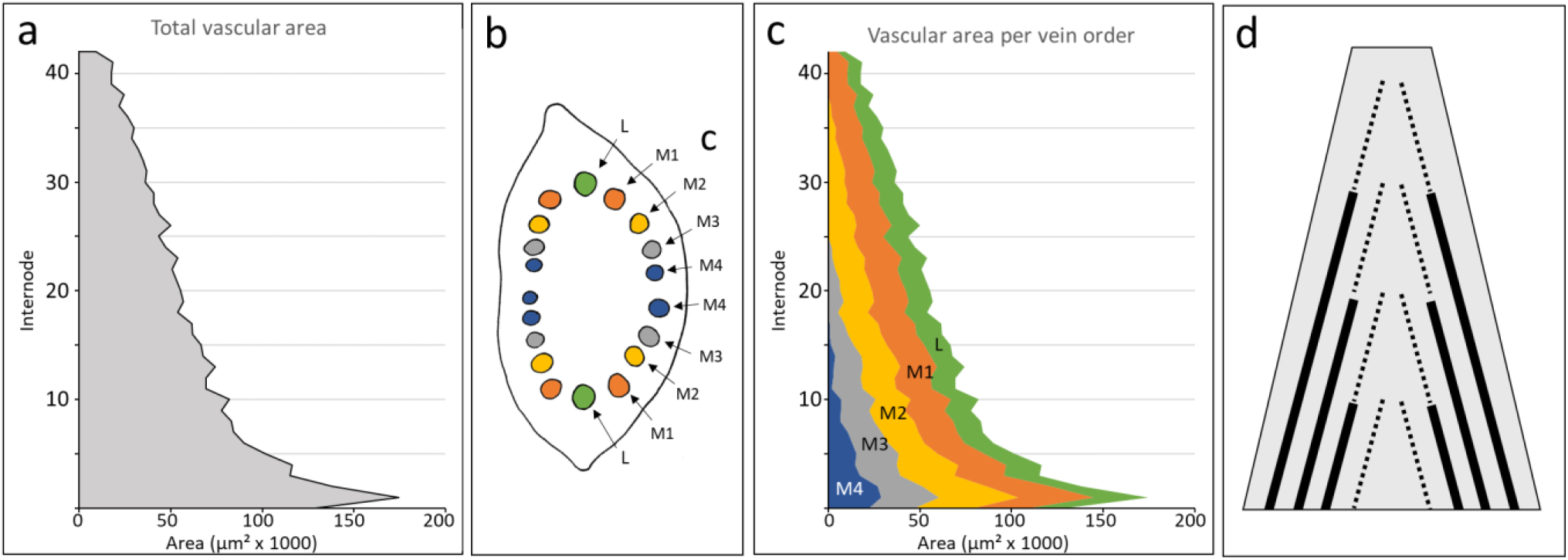
Contribution of individual vein orders to the overall vasculature along the rachis of a 43-node barley spike. (a) Total vascular area along the rachis. (b) Schematic overview of vascular organization in the 2^nd^ internode. Inwards of the lateral veins (L), the median veins are separated into 1^st^ (M1), 2^nd^ (M2), 3^rd^ (M3) and 4^th^ (M4) order median veins. (c) Differentiating the vascular area per vein order reveals the sequential diminishing and ultimate disappearing of 4^th^ (M4), 3^rd^ (M3) and 2^nd^ (M2) order median veins towards the apex. (d) Schematic presentation of vascular lay-out in the barley rachis with stable sized mature veins (solid lines) in the lateral periphery and a staggered array of diminishing immature veins (stippled lines) in the central area.

### Vein number decline along the rachis

The model for rachis vascular organization proposed in Fig. 6d predicts that rachis veins will ultimately fade into oblivion. If vein terminations are random, they should occur with similar regularity for both internodal (i.e. after passing a node, Fig. 7a) and intranodal (i.e. between two nodes, Fig. 7a). However, the summation of the positive and negative changes in the vein number along the rachis reveals a strong disparity between internodal (Fig. 7b) and intranodal vein number changes (Fig. 7c). Internodal changes start with a strong increase in vein number at the extreme base until internode 3, and then transforms into a consistent decline running at an average of close to 0.4 veins per internode (Fig. 7b). In contrast to this, intranodal changes are far less common, do not show an anomaly at the extreme base and, most significantly, are dominated by small vein number increases (Fig. 7c). The results thus suggest that most rachis veins ultimately end near or at a node. Furthermore, following the assumption that rachis veins do not merge or split within an internode (Kilby and Rymer, 1974), the small but consistent intranodal increase in vein number indicates that elements of the spikelet vasculature are entering and growing downward into the rachis.

**Fig. 7.**
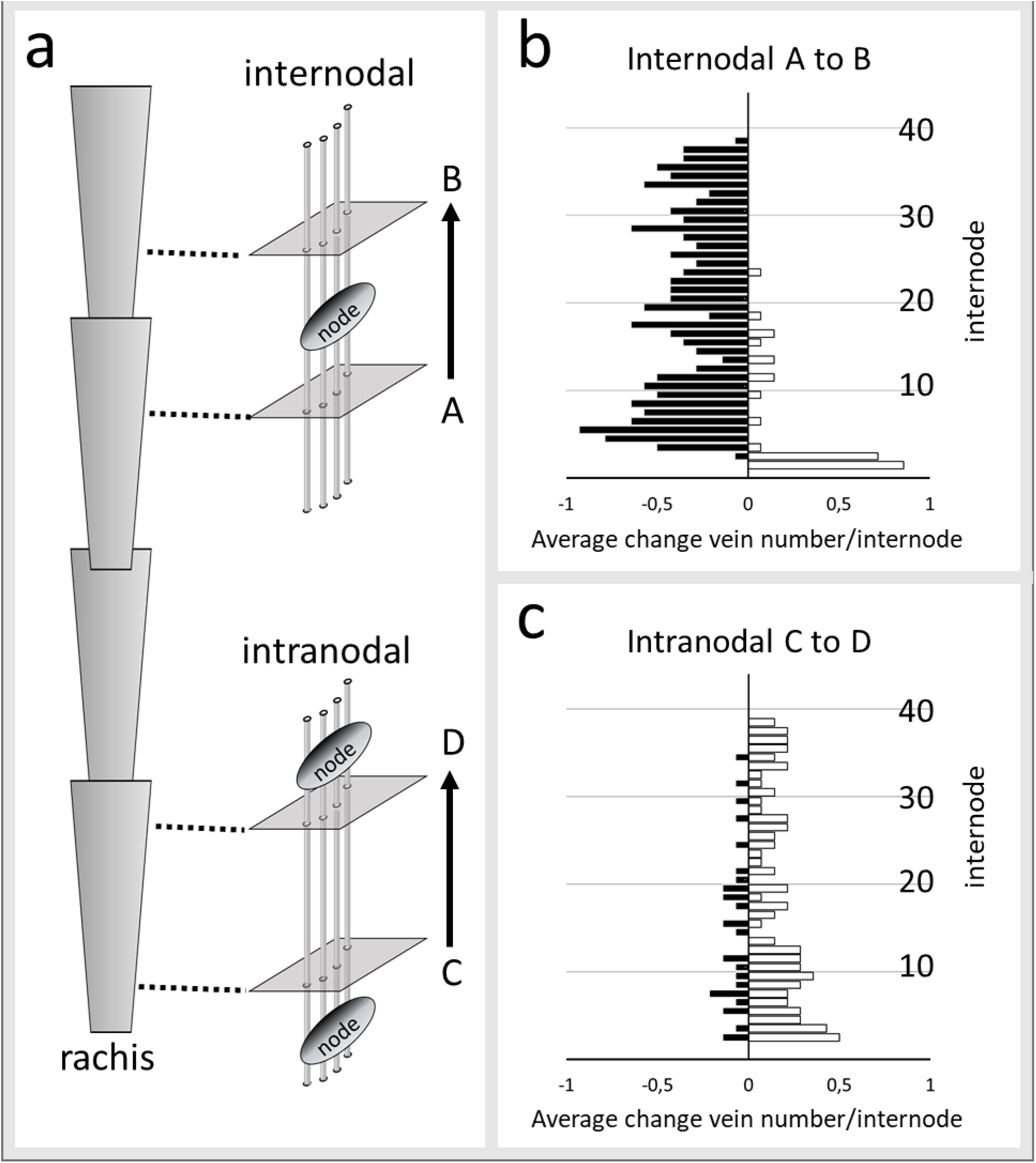
Internodal and intranodal changes in vein number along the rachis of mature spikes (average size 36,5 ± 2,9 nodes, n = 14). (a) Schematic view of measuring internodal and intranodal changes by serial internode sectioning. (b) Internodal changes start with a small zone of strong vein number increases (open bars) in the extreme base of the rachis. After a tipping point at internode 3, vein number decreases (solid black bars) become the dominant form of internodal vein number changes. (c) Intranodal vein number changes are less frequent than intranodal changes, lack an anomaly at the extreme base and are dominated by small but consistent vein number increases (open bars).

### Extra-numeral veins and rachis vasculature

Putative spikelet derived veins within the rachis expose themselves by what we here refer to as extra-numeral veins. This type of veins is usually distinctly smaller than their neighboring counterparts. Extra-numeral veins can occasionally be identified in the center of the rachis occupying an otherwise void space within the crescent row of median veins (Fig. 8a). The most exposed group of extra-numeral veins, however, is found in the periphery of the rachis, where they are located in between but slightly outside of the much larger lateral and first order median veins (Fig. 8b). The peripheral setting indicates that extra-numeral veins probably descends from the lateral spikelets. Analysis of dual consecutive internode sections suggests that upon entering the rachis extra-numeral veins continue in a basipetal direction while rapidly decreasing in size (Fig. 9a, b). Upon reaching the rachis extra-numeral veins may also fuse with the local rachis vasculature, cases of which were regularly detected in the central area of the rachis (Fig. 9c).

**Fig. 8.**
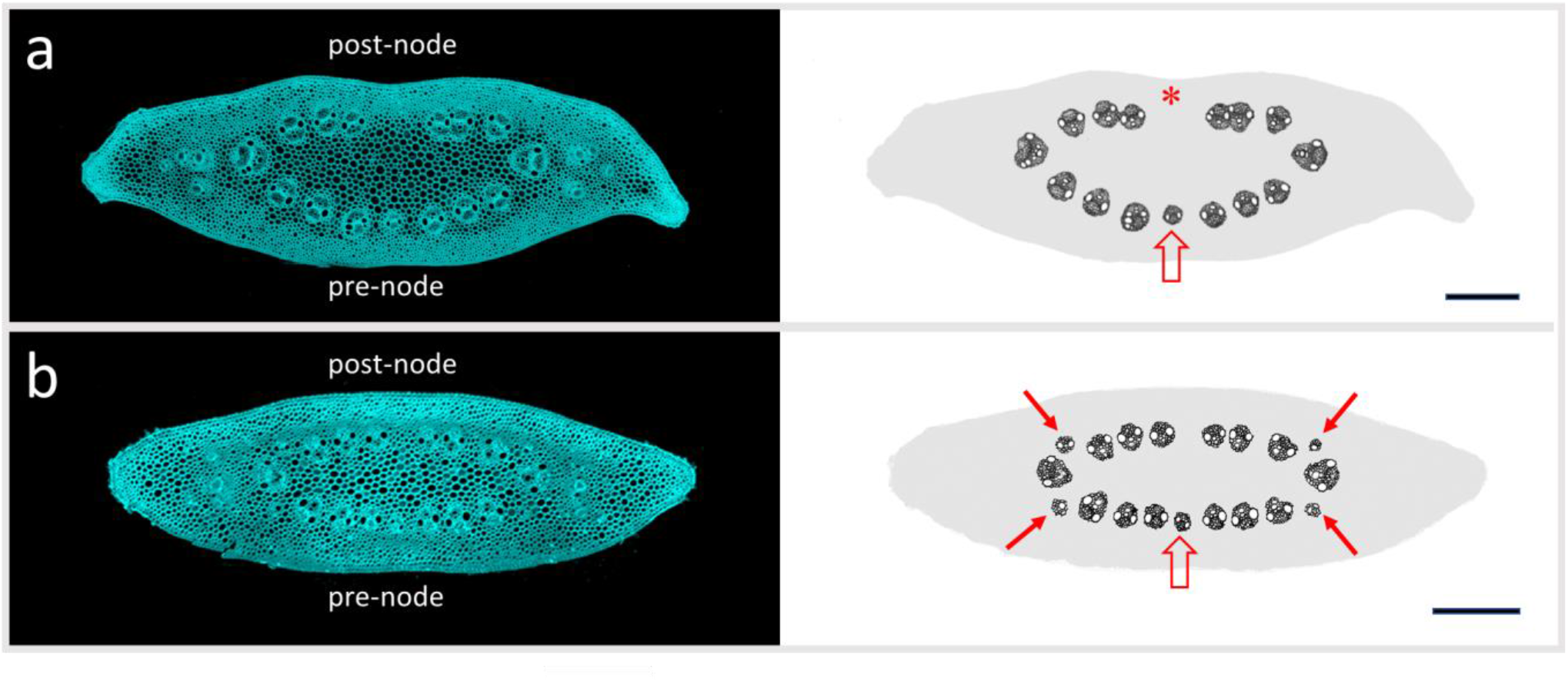
Presence of extra-numeral veins. (a) Internode no. 10 of a 36-node spike and schematic illustration of vascular distribution. The crescent row of median veins just apical of a node (post-node) reveals a conspicuous large empty area in the center (asterix). In the opposite row of veins which are about to reach a node (pre-node), the center is occupied by a notable small vein (open arrow). (b) Internode no. 14 of a 35-node spike and schematic illustration of vascular distribution. Veins that are distinctly smaller than their neighbors are found in the pre-node center (open arrow) and on all positions in between and slightly outside of the lateral and first order median veins (arrows). Bar = 200 µm.

**Fig. 9.**
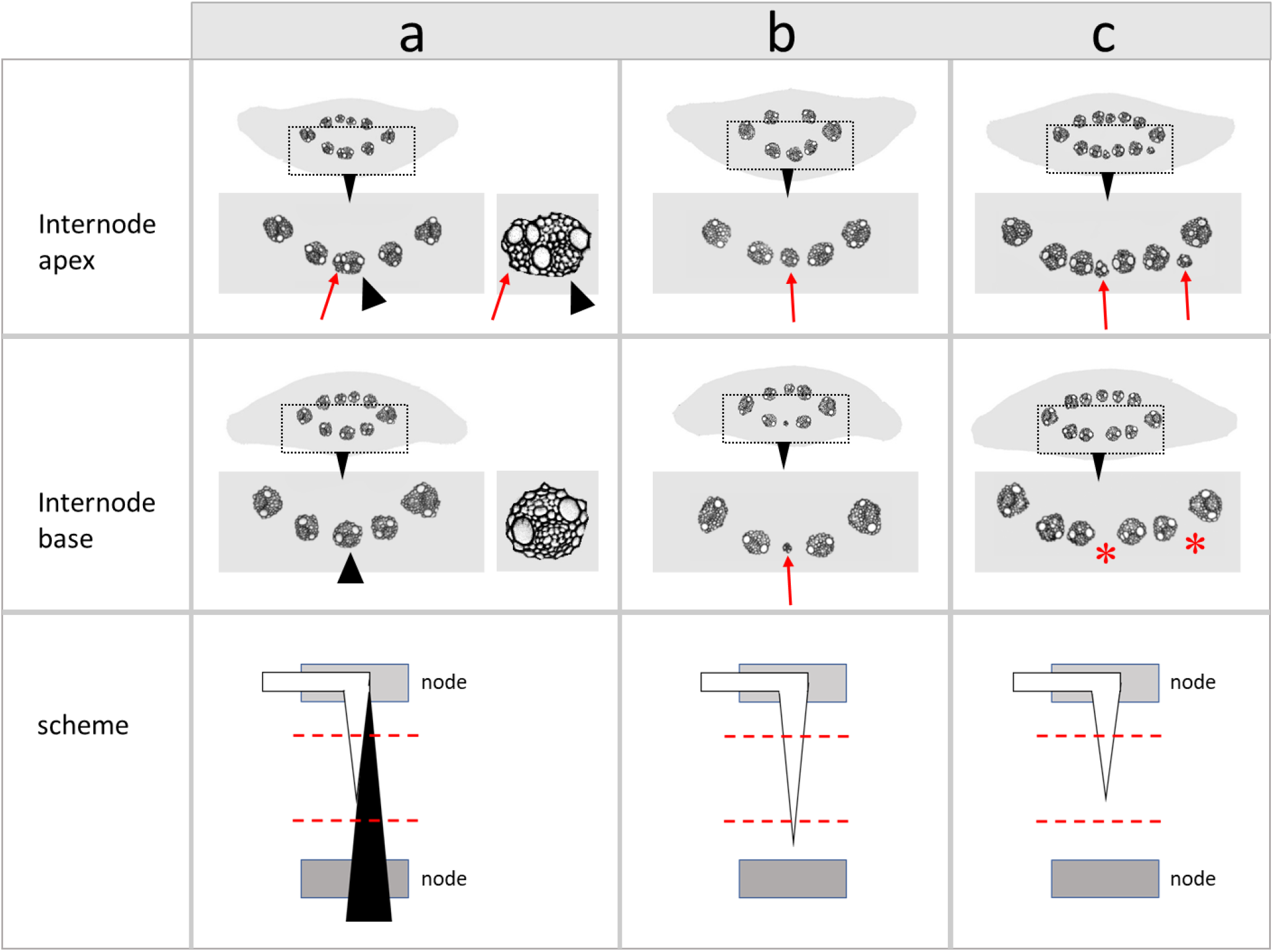
Identification of spikelet derived extra-numeral veins in paired internode sections. (a) spikelet vein (red arrows) fusing with a rachis vein (black arrowheads); (b) non-fusing spikelet vein basipetally decreasing in size; (c) non-fusing spikelet vein disappearing (asterix) before reaching next node. Schematic views show spikelet veins in white) and rachis veins in black veins while dashed red lines indicate the planes of sectioning. Spikelet veins predominant ly enter the central area of the rachis and only fuse with the immature rachis veins present there.

### Floret fertility and rachis vasculature

Transmutation of the two-rowed GP into the near-isogenic six-rowed GP*-vrs1* (Fig. 10a) triples the number of fertile spikelets per node without changing the number of spikelets per se. Despite this conversion, two-rowed GP (spike size 36,9 ± 0,9 nodes, n = 9) and six-rowed GP*-vrs1* (spike size 37,2 ± 0,7 nodes, n = 5) spikes with similar node number display virtually identical internode diameters along the rachis (Fig. 10b). Differences appear to reside solely in the vasculature. Including all veins into the comparison, then the rachis of the six-rowed GP*-vrs1* (14,8 ± 0,2 veins per internode) displays a small but significant increase (P > 0,001) in the vein number compared to the two-rowed GP (13,1 ± 0,9 veins per internode). Even when discounting for the large number of putatively spikelet-derived extra-numeral veins in between lateral and first order median veins (Fig, 11), the difference in the remaining vein number between two-rowed GP(12,8 ± 0,7 veins per internode) and six-rowed GP*-vrs1* (14,0 ± 0,1 veins per internode) remains significant (Fig. 10c). Far more striking though is the increase in total vascular area in the six-rowed transformant. Excluding the extra-numeral veins, the average total vascular area per internode in the six-rowed GP*-vrs1* is 26,2% larger than that in the two-rowed GP (Figs. 10e, 11). Analysis of laterals and first order median veins confirms that the transformation into a six-rowed phenotype is accompanied by a marked increase in overall vein diameter including an average 21.7% increase for lateral veins and 17,4% increase for 1^st^ order medial veins (Figs. 10e, 11). This identifies expansion of vascular diameter as the main cause for the rise in vascular area in the 6-row GP*-vrs1*.

**Fig. 10.**
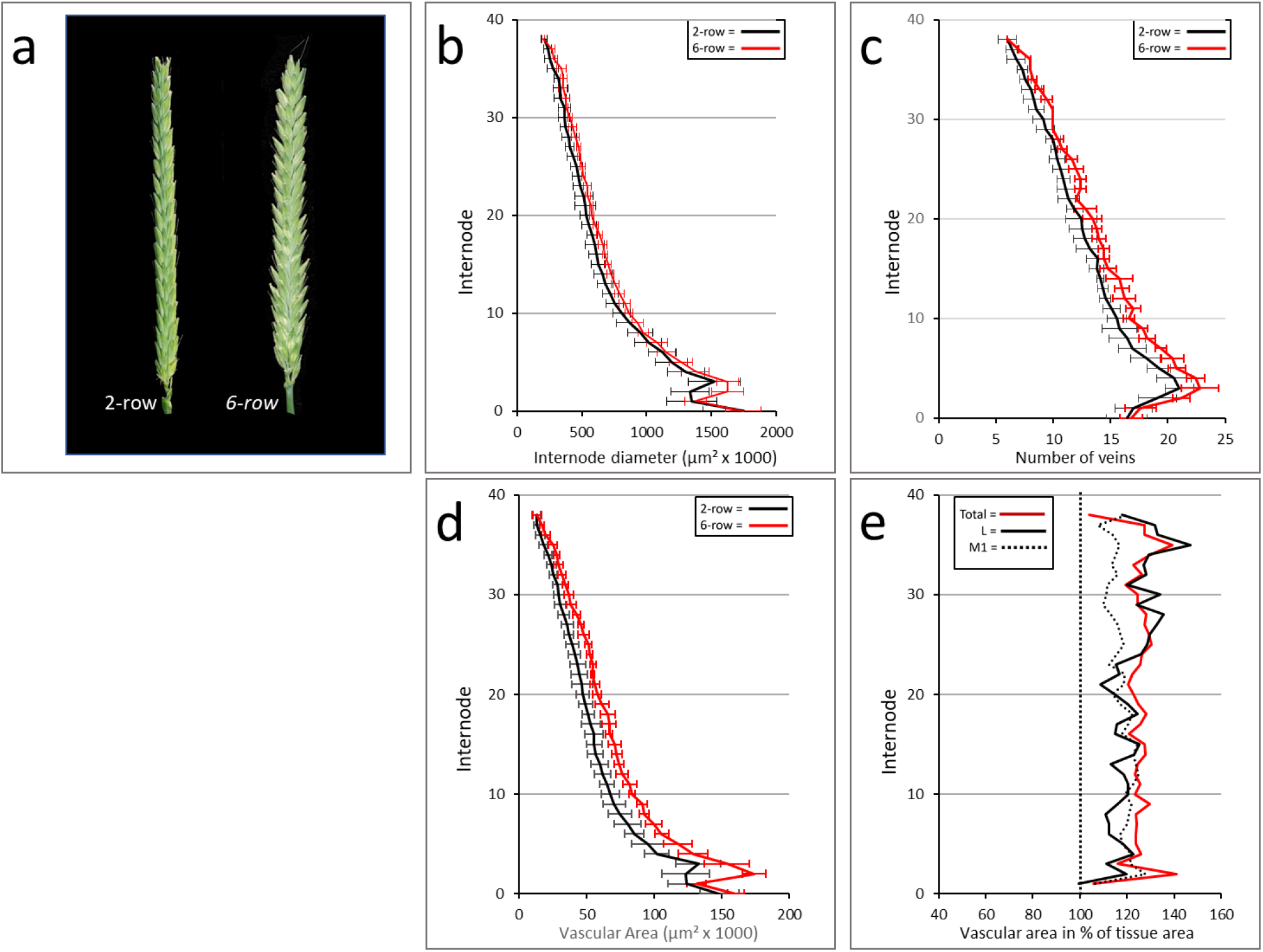
Floret fertility and rachis vasculature. (a) Spike phenotype of two-rowed GP and near isogenic six-rowed GP*-vrs1*. (b) two-rowed GP and six-rowed GP*-vrs1* n have similar rachis diameter. Even when extra-numeral veins are excluded, the six-rowed rachis contains more veins (c) and has a distinctly larger total vascular area (d) than the two-rowed rachis. (e) Against the normalized values for the two-rowed GP to 100% (straight vertical stippled line), the significant increase in total vascular area of the six-rowed conversion (solid red line) is caused by a general increase in vein size as shown for laterals (solid black line) and 1^st^ order median veins (stippled black line). For matters of clarity standard deviation bars were omitted. Bar in (a) = 1 cm.

**Fig. 11.**
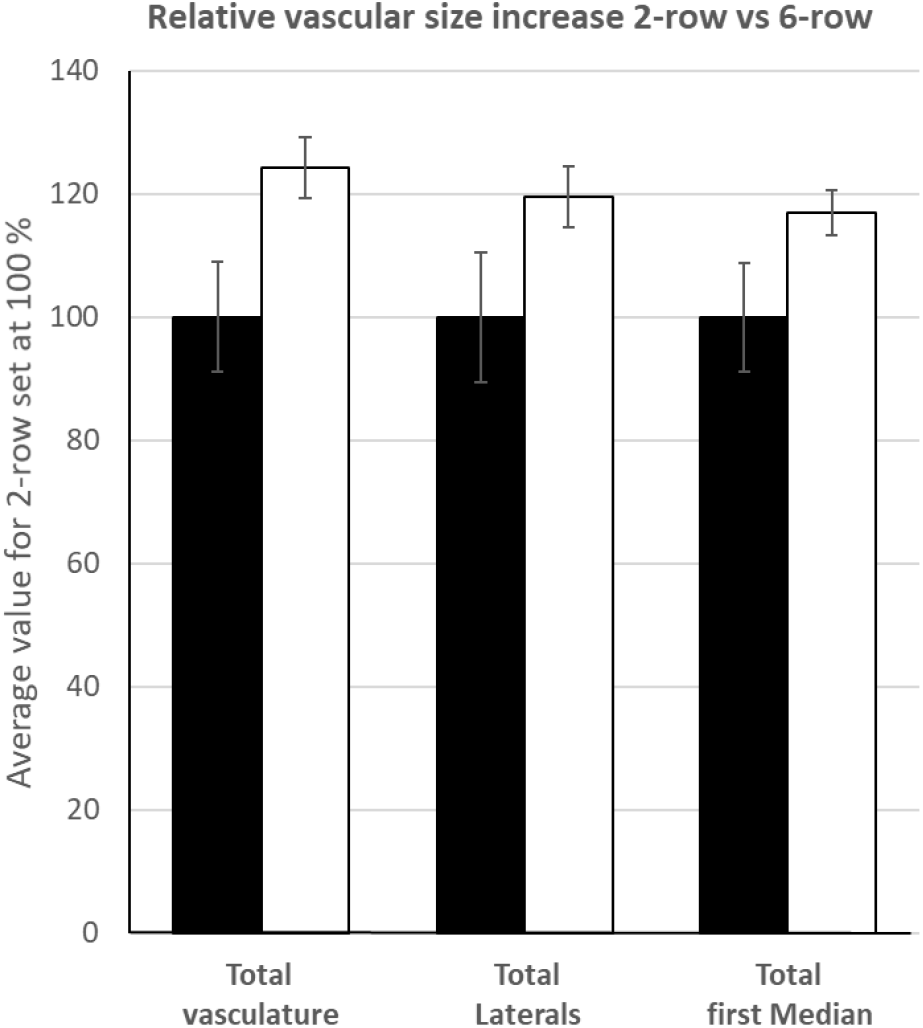
Increase in total vascular area and vascular dimensions of individual vein orders upon the conversion of the two-rowed GP (n=9) into the near isogenetic six-rowed GP-*vrs1* (n=5).

### Extra-numeral veins in two-rowed and six-rowed rachises

The subgroup of extra-numeral veins located between lateral and median vein in the two-rowed GP rachis appear to originate from the lateral spikelets. Serial internode section analysis of the six-rowed transformants supports this assumption. Furthermore, since there never was more than one extra-numeral vein between a lateral and first order median vein, it seems that each spikelet is connected to the rachis by just a singular extra-numeral vein. Like in the two-rowed GP, the number of extra-numeral veins in rachises of six-rowed GP*-vrs1* varies substantia l ly (Fig. 12a). Still, the distribution pattern along the rachis is similar between two-rowed GP and six-rowed GP*-vrs1* (Fig. 12b). Compared to the two-rowed GP, the six-rowed transformants contains on average about four times as many extra-numeral veins which are also substantia l ly larger (Fig. 12a-c). Furthermore, of the extra-numeral veins entering the peripheral part of rachis, only about 50% (21 out of 41) may reach the next basipetal node in the rachis of two-rowed GP, compared to more than 70% (60 out of 83) in the rachis of six-rowed GP*-vrs1*.

**Fig. 12.**
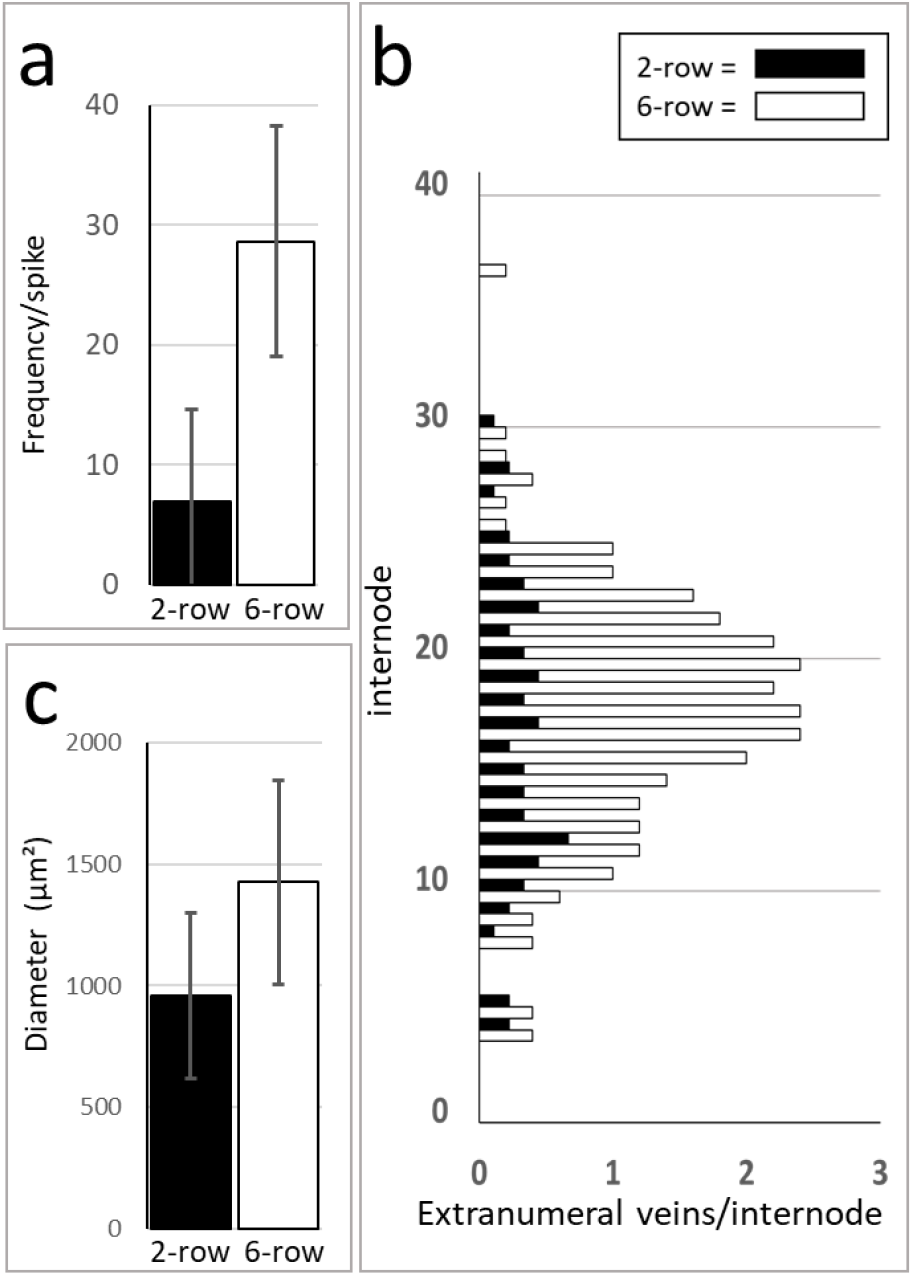
Peripheral extra-numeral veins in two-rowed GP (spike size36,9 ± 0,9 nodes, n = 9) and near isogenetic six-rowed GP*-vrs1* (spike size 37,2 ± 0,7 nodes, n = 5). (a) Frequency of extra-numeral veins in the six-rowed GP*-vrs1* is about four times higher than in the two-rowed GP. (b) Distribution analysis show that in both two-rowed GP and six-rowed GP*-vrs1* extra-numeral veins predominantly occur in the middle part of the rachis. (c) Average diameter of the six-rowed extra-numeral veins (n=142) is significantly larger (P>0.001) than that of the two-rowed extra-numeral veins (n = 62).

### Spikelet reduction and rachis vasculature

Barley plants exhibiting the *tst2.b* mutation undergo extended pre-anthesis tip degeneration resulting in a significantly shortened spike (Dahleen et al., 2007, Huang et al., 2023). To examine the effects on the rachis vasculature, spikes of cv. Bowman (spike size 25,3 ± 1.8 nodes, n = 11) were compared with those of the near isogenic mutant BW*-tst2.b* (spike size 14,2 ± 1.8 nodes, n =13) (Fig. 13a). Despite the severe reduction in spike size, the *tst2.b* mutation has no effect on either internode diameter or rachis vein number (Fig, 13c). Concerning total vascular area there are two main regions of disparities between wild-type and *tst2.b* exist (Figs. 13d): the first being a short region at the extreme base of the rachis while the second region starts and becomes progressively pronounced from about internode 8 onwards. These deviations become more evident when data for total vascular area, lateral veins and 1^st^ order median veins are compared. The results show that all vascular components of the *tst2.b* mutant are affected to the same extent in the extreme base and progressively so from internode 8 onwards (Fig. 13e). In the rachis section running from internode 3 to 8, however, total vascular area and diameter of individual veins do not differ between wild-type and mutant (Fig. 13e).

**Fig. 13.**
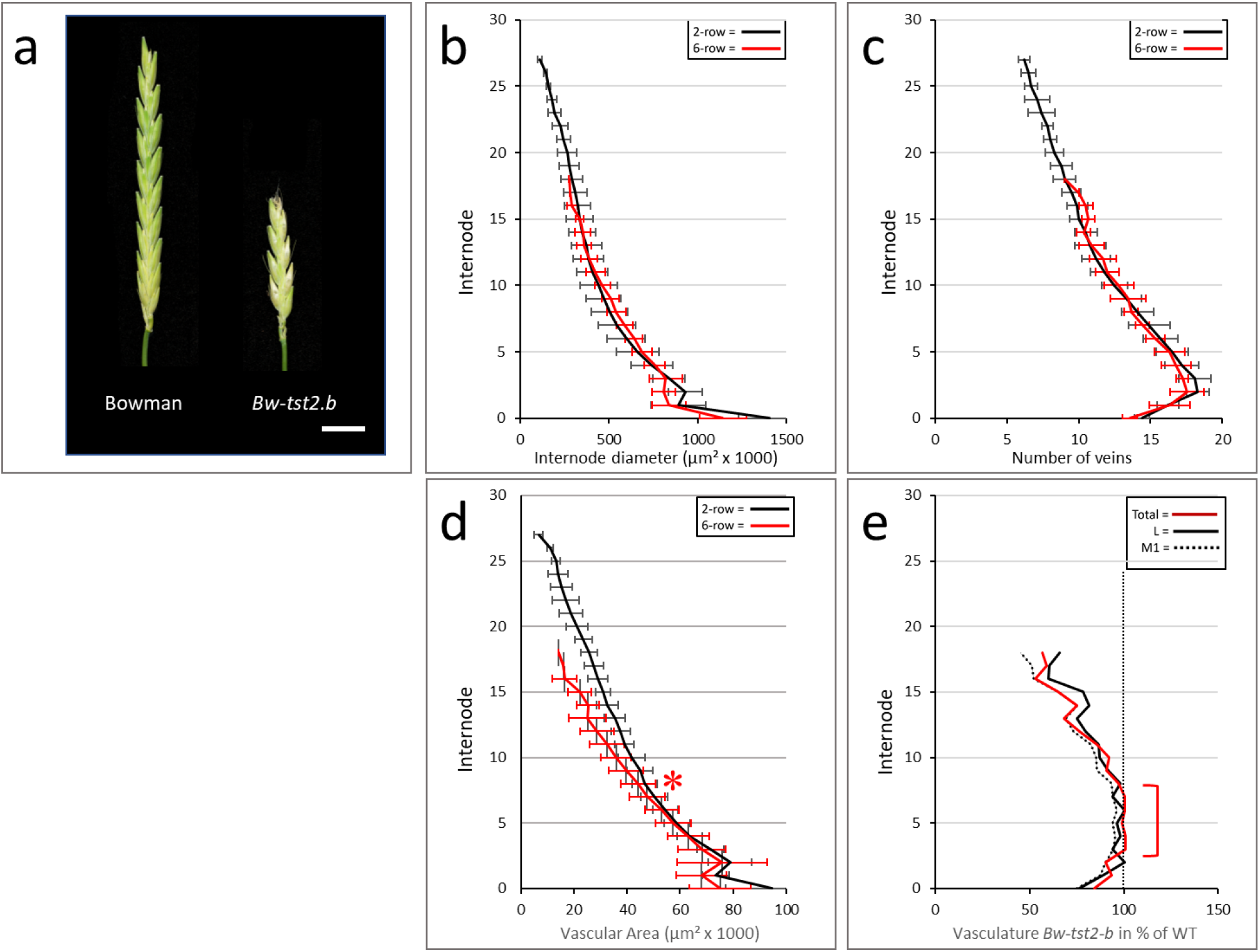
Effect of pre-anthesis tip degeneration on rachis vasculature. (a) Mature spikes of cv. Bowman and near isogenic mutant BW-*tst2.b*. Extensive pre-anthesis tip degeneration in the tst2-b mutant has no clear effect on internode surface area (b) and vascular number (c). (d) Towards the extreme base (arrow) and from about internode no. 8 onwards (asterix) total vascular area of the tst2-b is distinctly less than that in cv. Bowman. (e) Vascular inequalit ies between cv. Bowman and BW-*tst2.b* involve all vein classes to the same extent. Against the normalized values for cv. Bowman to 100% (straight vertical stippled line), in BW-*tst2.b* total vascular area (red line) as well as diameter of laterals (black line) and 1^st^ order median veins (stippled black line) are significantly reduced at the extreme base, nearly similar between internode 3 to 8 (brace), and become progressively smaller again beyond internode 8. For matters of clarity standard deviation bars were omitted in (e). Bar in (a) = 1 cm

### Rachis parameters in wild barley

A comparative analysis of wild barley accession HID003 (average spike size 22,0 ± 1.4 nodes, n = 8) shows that its vascular lay-out is similar to that of the barley two-rowed GP and Bowman with the number of main vascular bundles reaching a peak around internode 3 (compare Figs. 2c, 13d, 14d). Concerning the parameters rachis diameter and vascular area there are differe nces though. Whereas in two-rowed GP and Bowman these parameters show more or less distinct two-stepped declines (Fig. 2b-c, 13b-c), the declines in HID003 are near to linear (Fig. 14b-c) in which especially the vascular area decline resembles the profile seen in BW-*tst2.b* (Fig. 14b, c). The dynamics of individual vein orders emphasizes basic differences between the barley cultivars and wild barley accession HID003, while also accentuating the similarities between the latter and BW*-tst2.b* (Fig. 15). In the extreme base of the rachis vascular bundles in two-rowed GP and Bowman show steep acropetal declines in size (arrows in Fig. 15a-b) whiles those in HID003 and BW*-tst2b* first increase to a more or less pronounced maximum value at internode 2 or 3 before declining (Asterix in Fig. 15c-d). Overall, the declines in vascular size along the rachis describes faint S-shaped curves in two-rowed GP and Bowman (Fig. 15a-b) but are more linear in HID003 and BW*-tst2.b* (Fig. 15c-d).

**Fig. 14.**
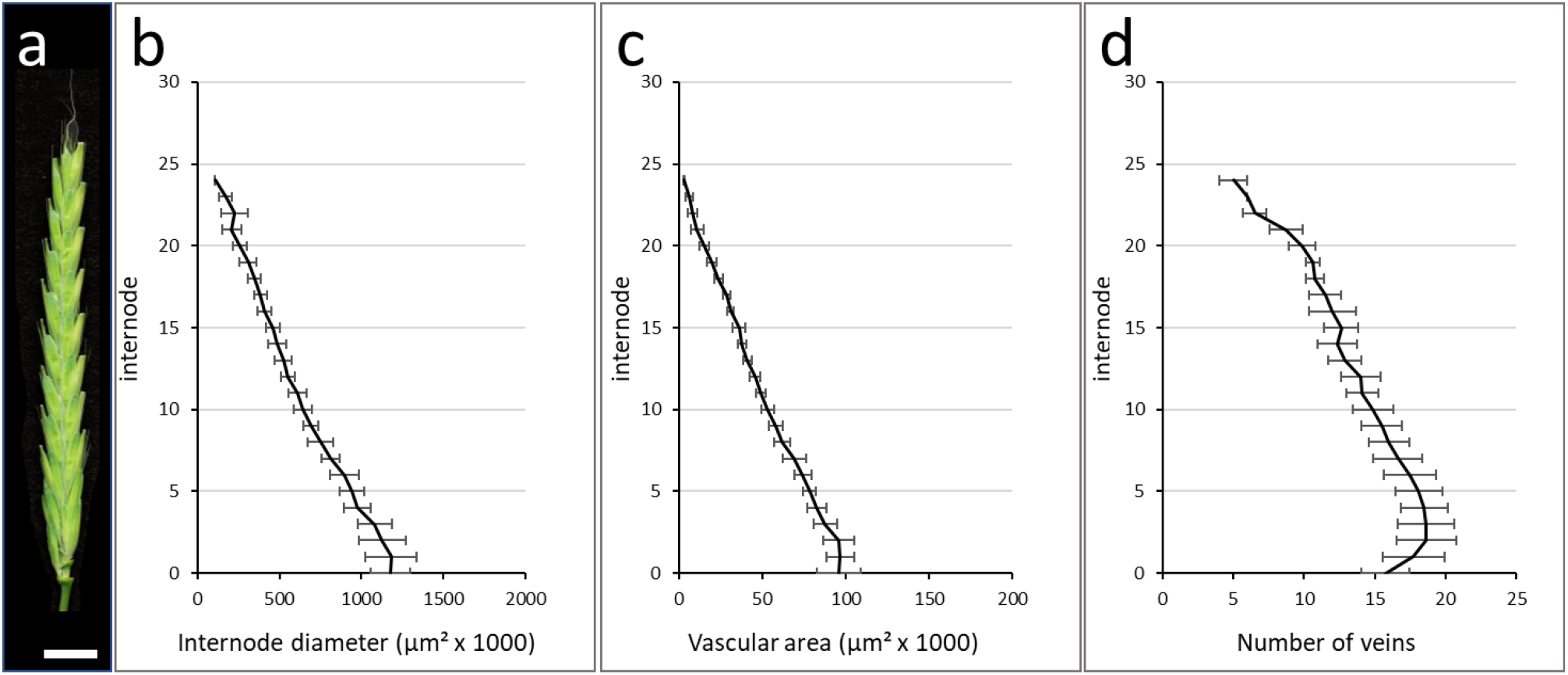
Dynamics of internode parameters along the rachis of wild barley (*Hordeum spontaneum*) acc. HID003. Data was collected from mature spikes between 20 to 24 nodes in size (average 22.0 ± 1.4 nodes, n = 8). (a) mature spike. With the possible exceptions of the extreme base, both internode diameter (b) and vascular area (c) display near linear declines along the rachis. (d) Vascular number reaches a maximum value around internode 3 before gradually declining towards the tip. Bar in (a) = 1 cm.

**Fig. 15.**
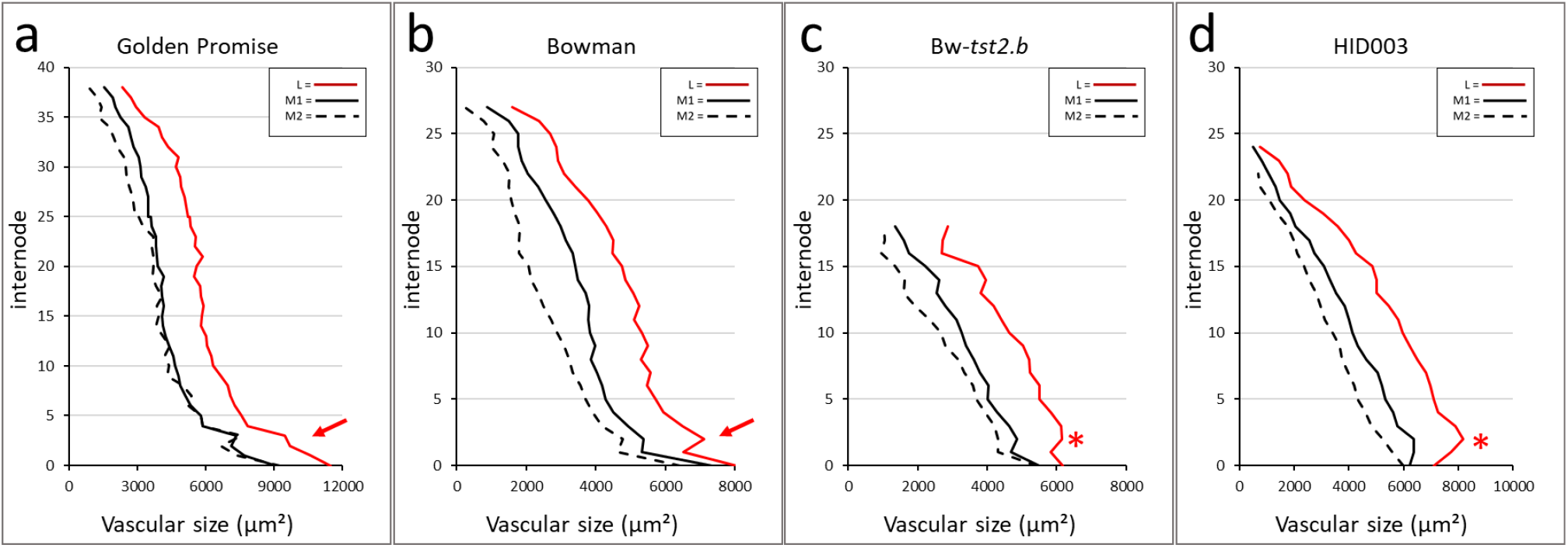
Comparative analysis of individual vein order dynamics along the rachis of two-rowed GP (a), cv. Bowman (b), BW-tst2.b (c), and wild barley accession HID003 (d). For matters of clarity standard variations were omitted. At the base of the rachis vascular bundles in cvs. Golden Promise and Bowman steeply decline in size (arrows in a-b) whiles vascular bundles in mutant BW*-tst2.b* and wild barley accession HID003 first reach maximum values at internode 2 or 3 (asterix in c and d) before declining. L = lateral veins, M1 = 1^st^ order median vein, M2 = 2^nd^ order median vein.

## Discussion

Barley is considered to be a sink-limited cereal in which assimilate supply exceeds the storage capacity of the grains (Bingham et al., 2007). Also transport capacity of the rachis vasculature is in excess of the demands during grain filling (Yang & Zhang, 2006). This apparent absence of bottle-neck features may explain why the barley rachis vasculature has received less research in the past near half century. As a follow up on the pivotal study by Kirby and Rymer (1975) on barley rachis vasculature, the main aim of the present work was to clarify the relationship between vasculature and spike size, define vascular dynamics and provide an improved model for overall vascular organization within the rachis.

### Vascular parameters and spike size

The transition from a shoot to a spike phenotype takes several internodes to materialize [**supplementary information – video**]. This also accounts for the vascular organizat io n. Although often implied (Percival, 1921, Whingwiri et al., 1981), features like vascular number and size do not go through a gradual linear decline along the rachis (Fig. 2). Most conspicuous is that the number of main vascular strands displays an absolute maximum around internode three to four irrespective of the source material used (Fig. 2e, 11c, 12c, 13c). This may unveil the vein initiation site within the early rachis from which veins develop both acropetally into the apex and basipetally into the culm. Though Kirby and Rymer (1975) situated this site between spikelet primordium six and ten, observations by Pizzolato (1997) suggest a much more basal location.

The morphogenesis of the barley rachis vasculature appears to be the result of two distinct processes: initiation and processing. The basic lay-out of the spike including that of the rachis vasculature, is concluded by the time of maximum yield potential while differentiation of the initiated organs is finished by anthesis (Waddington et al., 1983, Thirulogachandar and Schnurbusch, 2021). Using mature spikes at the time of grain filling we accordingly found vein number only weakly correlated to spike size (Fig. 4c) while internode diameter and vascular area were strongly correlated (Fig. 4a-b). Correlations between vascular parameters and internode diameter, however, appeared very stable and unrelated to spike size (Fig. 5a-c) for which reason the factor internode diameter can be used for a comparative analysis of transport capacity between individual spikes (Rosell et al., 2017).

### Vascular organization and transport efficiency

Transport resistance is directly proportional to the length of a vessel, but inversely proportional to the radius to the fourth power (Tyree and Ewers, 1991, Cruiziat et al., 2002, Chiavarria and Pessoa dos Santos, 2012). This makes vessel diameter a central factor in transport efficiency with small changes having strong effects (Rosell et al., 2017). The strong correlation between rachis size and vascular size (Figs. 4b) confirms that transport capacity of the rachis is modelled mainly by vein size. This emphasis on transport efficiency is also reflected in the organiza t ion and dynamics of lateral and median vein orders (Fig. 6). During spike development new veins are initiated inwards of already existing ones (Kirby and Rhymer, 1975). Measurements of individual vein orders (Fig. 6c) not only suggest a near uniform size for all fully developed median veins (Fig. 6), but indicate that the initiation of a new order of veins coincides with the previous order reaching maturity (Figs. 6c).

It is tempting to speculate that the termination of the inflorescence meristem around the awn primordium stage not only determines the maximum yield potential (Thirulogachandar and Schnurbusch, 2021), but also puts a hold on any further vascular development, freezing the vascular bundles of the rachis in their respective stages of initiation. As such the central area of the barley rachis appears to be occupied by a staggered array of small, putative immature veins (Fig. 6d). Since marks of immaturity include the absence or incompleteness of a bundle sheath (Esau, 1943) the central rachis may thus be a stockpile of potentially leaky veins suited to provide a continuous supply of nutrients and metabolites along the central axis of the rachis. In contrast, the large mature veins in the rachis periphery may serve a transport role since their constant diameter should mitigate long distance transport resistance and facilitate apical supply. Observations on the uptake and distribution of labeled sucrose in the barley spike by Melkus et al., (2011) showed signals accumulating in the central area of the rachis, i.e. at the spikelet attachment site. This supports our hypothesis of a spatial separation between bulk transport taking place in the rachis periphery while local accumulation and delivery to the spikelets occurs in the center of the rachis.

### Interaction between spikelet and rachis vasculature

Once veins have reached the center of the rachis they start to diminish in size (Fig. 6) and ultimately end near or at a node (Fig. 7). The center of the rachis is also one of two locations where presumed spikelet-derived extra-numeral veins enter, the other location being in the periphery between lateral and 1^st^ order median vein (Fig. 8). Our observations on two-rowed and six-rowed barley suggest that from each spikelet only a singular vein extends towards and into the rachis. Once in the rachis, these veins, in analogy with leaf veins, probably extend basipetally and ultimately fuse with the rachis vasculature (Hitch and Sharman, 1968, Nelson and Dengler, 1997, Pizzolato, 1997). Through fusion events such as shown in Fig. 9a, the declining veins occupying the center of the rachis may over the course of several nodes be connected to multiple spikelet-derived veins. The latter may eventually “cap off” the rachis vein thus giving the impression of these veins ending at a node (Fig. 7).

Not all extra-numeral veins will merge with the rachis vasculature which is the very reason spikelet-derived veins become visible in the first place (Figs. 7c, 9b, c). This failure to fuse is especially evident in the rachis periphery and is particularly prominent in the six-rowed GP-*vrs1* line (Fig. 11). Given that most fusion events observed took place in the center of the rachis, it is tempting to speculate that this may reflect an overall inability of spikelet veins to merge with the large mature veins of the rachis periphery whereas they might readily fuse with the small immature veins occupying the rachis center. The high incidence with which in the six-rowed GP-*vrs1* veins originating from the lateral spikelets do not fuse with the rachis vasculature also questions the relevance of vascular continuity between spikelet and rachis vasculature. Kirby and Rymer (1974) remarked that when rachis veins transverse a node, i.e. the main place for nutrient exchange, it can be near to impossible to distinguish these veins from the surrounding parenchymal tissue rich in putative transfer cells. The facultative character of vascular continuity suggests that both vascular systems may fulfil their roles independently, the rachis veins delivering nutrients to the node and the spikelet veins collecting nutrients from the node. A fusion between the two vascular systems near or at the node may then be no more than a non-essential coincidence. Although vascular discontinuity also exists in the floral axis (Zee and O’Brien, 1970, Kirby and Rymer, 1975), it will require a 3D reconstruction to elucidate the true interaction between rachis and spikelet vasculature.

### Floret fertility and rachis vasculature

<u>The events that differentiate between a two-rowed GP and a six-rowed</u> GP*-vrs1* <u>establish themselves after</u> the stage of maximum yield potential (Thirulogachandar et al., 2021), i.e. after the basic lay-out of the rachis, including its vasculature, has been concluded. In accordance with this the transmutation of the two-rowed GP into a six-rowed GP*-vrs1*, did not affect the diameter of the rachis per se, and the number of rachis veins only slightly (Figs. 10b-c). The most prominent consequences were the increase in the number of extra-numeral veins in the lateral areas of the rachis (Fig. 11) and the significant rise in total vascular area (Fig. 10d), predominantly caused by an increase in vein diameter (Fig. 12e, Table 2). Concerning the fixed ratio between the dimensions of vascular bundles and metaxylem vessels (Fig. 4), a 20% increase in diameter should according to the Hagen-Posseuille equation (V/t = (r4 × π × ΔP) / (8 × η × l,) suffice to increase xylem transport capacity of the rachis by a factor of two. It thus seems that spikelet/floret fertility in the *vrs1* mutant has a quantitative effect on transport capacity along the whole rachis. Since Sakuma et al. (2013) discovered *Vrs1* to be localized in the vasculature of the developing rachis, this gene is likely to have role in vascular development. To examine possible consequences of the opposite, i.e. a reduction in spikelet/floret fertility, the cv. Bowman was compared with its near isogenic line BW*-tst2.b*, showing extended pre-anthesis tip degeneration (Fig. 12a) due to a mutation in a vascular-expressed *CCT MOTIF FAMILY4* (*HvCMF4*) gene (Huang et al., 2023). This mutation reveals itself after maximum yield potential (Huang et al., 2023) and mainly affects the dimensions of the veins while internode diameter and vein number itself remain unaltered (Fig. 12b-c). The vascular modifications involve all veins in a similar fashion (Fig. 12e). Far from being uniform along the rachis, the effect was pronounced in the extreme base, near to absent between internodes 3 to 8 and from then on becoming progressively stronger again towards the apex (Fig. 12d-e). In the last internode, average vein size had decreased by about 40% (Fig. 12e), reducing the xylem transport capacity to a mere 10% of that in the wild-type. This not only supports observations by Huang et al. (2023) implying that an altered vasculature predates pre-anthesis tip degeneration, but also underlines the importance of histological data for our understanding of the physiological functions of the barley spike.

### Barley domestication and rachis vasculature

In all barley plants studied here, vascular number dynamics always revealed a peak value slightly above the rachis base identifying this as the likely vein initiation site. A main difference between barley cultivars and both wild barley acc. HID003 and mutant BW-*tst2.b* is that in the latter two the site of maximum vein number overlaps with that of maximum vein size (Figs. 13, 14). As a consequence of this maximum transport capacity is reached well above the base with the latter thus becoming a potential bottle-neck. It seems conceivable that the domestication of barley may also have favored genotypes without this bottleneck. As work by Huang et al (2023) has shown, single alleles can be responsible for increasing the vascular dimensions in both the base and apical region of a spike thus greatly affecting spike yield.

## Conclusion

The morphogenesis and organization of the rachis vasculature is a multistep process. In the first phase the basic lay-out is established featuring the ultimate number of main rachis veins (Waddington et al., 1983, Thirulogachandar and Schnurbusch, 2021). This is followed by a second phase in which the dimensions of the rachis veins, and with it, the transport capacity of the rachis as a whole are adjusted to accommodate for future demands. In this, one of the determining factors appears to be spikelet fertility. Within the rachis vasculature we can differentiate between mature veins in the lateral periphery in charge of long distance transport, and a staggered array of immature veins in the center responsible for local spikelet supply (Fig. 9, 15). The center of the rachis also serves as the main port of entry for spikelet-derived veins, the other positions are located in the rachis periphery (Figs. 10, 11). The frequently observed failure of spikelet-derived veins to merge with the rachis vasculature, indicates that vascular continuity between spikelets and rachis may be a non-essential feature. Results obtained for wild barley acc. HID003 suggests that the domestication of barley may have favored the selection for genotypes with reduced vascular bottle-necks.

## Supplementary information – video

Serial transversal internode sectioning through the rachis of a barley spike cv. Golden Promise. Starting at the peduncle 2 cm underneath the rachis proper, all subsequent sections were made at about 1/3 of the internodal height. The sections reveal the changing phenotype of the rachis next to the altering organization of the rachis vasculature and the progressive decline in vascular number towards the apex. Solid green ovals indicate the position of the next acropetal fertile spikelet attachment site. The open green oval in last internode 36 indicates the start of the apical abortion zone.

## ACKNOWLEDGEMENTS

We thank Enk Geyer, Daniela Feldmann and Anke Mueller for assistances with greenhouse plant care and Dr. Armin Meister for help with statistical analysis.

## Reference

1. Alqudah AM, Schnurbusch T. 2014. Awn primordium to tipping is the most decisive developmental phase for spikelet survival in barley. Functional Plant Biology 41: 424–436.

2. Appleyard M, Kirby E, Fellowes G. 1982. Relationships between the duration of phases in the pre-anthesis life cycle of spring barley. Australian Journal of Agricultural Research 33: 917–925.

3. Bingham I, Blake J, Foulkes MJ, Spink J. 2007. Is barley yield in the UK sink limited? II. Factors affecting potential grain size. Field Crop research 101: 212–220.

4. Boussora F, Allam M, Guasmi F, et al. 2019. Spike developmental stages and ABA role in spikelet primordia abortion contribute to the final yield in barley (Hordeum vulgare L.). Botanical Studies 60: doi: 10.1186/s40529-019-0261-2.

5. Bremner PM. 1972. Accumulation of dry matter and nitrogen by grains in different positions of the wheat spike as influenced by shading and defoliation. Australian Journal of Biological Sciences 25: 657–668.

6. Bremner PM, Rawson HM. 1978. The weights of individual grains of the wheat spike in relation to their potential, the supply of assimilate and interaction between grains. Functional Plant Biology 5: 61–72.

7. Budhagatapalli N, Schedel S, Gurushidze M, et al. 2016. A simple test for the cleavage activity of customized endonucleases in plants. Plant Methods 12: 18, doi:10.1186/s13007-016-0118-6.

8. Cruiziat P, Cochard H, Ameglio T. 2002. Hydraulic architecture of trees: main concepts and results. Annals of Forest Science 59: 723–752.

9. Dahleen L, Franckowiak JD, Lundqvist U. 2007. Descriptions of barley genetic stocks for 2007. Barley Genetics Newsletter 37: 154–187.

10. Esau K. 1943. Ontogeny of the vascular bundle in *Zea Mays*. Hilgardia 15: 325–368.

11. Evans LT, Dunstone RL, Rawson HM, Williams RF. 1970. The phloem of the wheat stem in relation to requirements for assimilate by the spike. Australian Journal of Biological Sciences 23: 743–52.

12. Hanif M, Langer RHM. 1972. The vascular system of the spikelet in wheat (Triticum aestivum). Annals of Botany 36: 721–727.

13. Hensel G, Kastner C, Oleszczuk S, Riechen J, Kumlehn J. 2009. Agrobacterium-mediated gene transfer to cereal crop plants: current protocols for barley, wheat, triticale, and maize. International Journal of Plant Genomics. 2009:2009:835608. doi:10.1155/2009/835608.

14. Huang Y, Kamal R, Shanmugaraj N, et al. 2023. A molecular framework for grain number determination in barley. Science Advances 9: eadd0324. doi:10.1126/sciadv.add0324

15. Kato T. 2004. Effect of spikelet removal on the grain filling of Akenohoshi, a rice cultivar with numerous spikelets in a panicle. Journal of Agricultural Science 142: 177–181.

16. Kirby EJM, Faris DG. 1970. Plant population induced growzh correlations in the barley shoot and possible hormonal mechanisms. Journal of Experimental Botany 21: 787–798.

17. Kirby EJM, Rymer JL. 1974. Development of the vascular system in the ear of barley. Annals of Botany 38: 565–573.

18. Kirby EJM, Rymer JL. 1975. The vascular anatomy of the barley spikelet. Annals of Botany 39: 205–211.

19. Koppolu R, Schnurbusch T. 2019. Developmental pathways for shaping spike inflorescence architecture in barley and wheat. Journal of Integrative Plant Biology 61: 278–295.

20. Ma YZ, Mackown CT, Van Sanford DA. 1996. Differential effects of partial spikelet removal and defoliation on kernel growth and assimilate partitioning among wheat cultivars. Field Crop Research 47: 201–209.

21. Melkus G, Rolletschek H, Fuchs J, et al. 2011. Dynamic 13C/1H NMR imaging uncovers sugar allocation in the living seed. Plant Biotechnological Journal 9: 1022–1037

22. Nelson T, Dengler N. 1997. Leaf vascular pattern formation. Plant Cell 9: 1121–1135.

23. Percival J. 1921. The wheat plant, 463 pp, Duckworth and co., London.

24. Pizzolato TD. 1997. Procambial initiation for the vascular system in the spike of wheat. International Journal of Plant Sciences 158: 121–131.

25. Rosell JA, Olson ME, Anfodillo T. 2017. Scaling of xylem vessel diameter with plant size: causes, predictions, and outstanding questions. Current Forestry Reports 3: 46–59.

26. Sakuma S, Pourkheirandish M, Hensel G, et al. 2013. Divergence of expression pattern contributed to neofunctionalization of duplicated HD-Zip I transcription factor in barley. New Phytologist 197: 939–948.

27. Schindelin J, Arganda-Carreras I, Frise E, et al. 2012. Fiji: an open-source platform for biological-image analysis. Nature Methods 9: 676–682.

28. Thirulogachandar V, Schnurbusch T. 2021. ‘Spikelet stop’ determines the maximum yield potential stage in barley. Journal of Experimental Botany 72: 7743–7753.

29. Thirulogachandar V, Govind G, Hensel G, et al. 2023. Barley SIX-ROWED SPIKE 1 regulates spikelet development independently of its evolutionarily conserved paralog, HOMEOBOX2. submitted to JXB, 10.1101/2021.11.08.467769

30. Waddington SR, Cartwright PM, Wall PC. 1983. A quantitative scale of spike initial and pistil development in barley and wheat. Annals of Botany 51: 119–130.

31. Whingwiri EE, Kuo J, Stern WR. 1981. The vascular system in wheat. Annals of Botany 48: 189–202.

32. Whingwiri EE, Kemp DR. 1980. Spikelet development and grain yield of the wheat spike in response to applied nitrogen. Australian Journal of Agricultural Sciences 31: 637–647.

33. Yang J, Zhang J. 2006. Grain filling of cereals under soil drying. New Phytologist 169: 223–236.

34. You C, Zhu H, Xu B, et al. 2016. Effect of Removing Superior Spikelets on Grain Filling of Inferior Spikelets in Rice. Frontiers of Plant Sciences 7: 1161.

35. Zee SY, O’Brien TP. 1970. A special type of tracheary element associated with “xylem discontinuity” in the floral axis of wheat. Australian Journal of Botany 23: 783–791.

